# The lytic transglycosylase MltA participates in turnover of septal peptidoglycan in *Escherichia coli*

**DOI:** 10.64898/2026.05.07.723478

**Authors:** Atsushi Yahashiri, Gabriela M. Kaus, David S. Weiss

## Abstract

Daughter cell separation in *Escherichia coli* is driven primarily by two classes of peptidoglycan (PG) hydrolases that work in tandem: *N*-acetylmuramoyl–L-alanine amidases that strip stem peptides from the PG glycan backbone and lytic transglycosylases (LTs) that break down the PG glycan backbone. Although the relevant amidases have been known for years, which of *E. coli’s* eight LTs contribute to this process is less clear. Because the amidases process PG first, the relevant LTs must utilize peptide-free or “denuded” glycan substrates (dnGs). MltA is one of the few *E. coli* LTs that can break down peptide-free PG glycans *in vitro*, but its precise physiological roles are not known. Here we show MltA localizes to the division site in constricting *E. coli* cells and cells lacking MltA accumulated dnGs in septal PG. We found that MltA binds to the anhydroMurNAc ends of glycan chains, which raises the possibility that these structures are enriched in septal PG. Nevertheless, as reported previously, deletion of *mltA* does not impair daughter cell separation sufficiently to cause a chaining phenotype. Overall, our findings demonstrate that MltA is a physiologically relevant peptidoglycan hydrolase for cell division in *E. coli*.

**IMPORTANCE:** How bacteria coordinate synthesis and cleavage of septal peptidoglycan remains poorly understood, in part because some of the relevant enzymes have yet to be identified. Here we show that the *E. coli* lytic transglycosylase MltA is involved in cleaving septal peptidoglycan. Besides elucidating a physiological role for MltA, our work brings the field a step closer to identifying all of the proteins involved in cell division in an important model organism.

## INTRODUCTION

Most bacteria are surrounded by a wall of peptidoglycan (PG; also called murein) that is the ultimate target of many important antibiotics (1–3). PG consists of glycan strands connected by peptide crosslinks to create a net-like sacculus that confers cell shape and helps to prevent turgor pressure from rupturing the membrane (4, 5). The glycans are built from a repeating disaccharide of *N*-acetylglucosamine (GlcNAc) and *N*-acetylmuramic acid (MurNAc). The peptides used for crosslinking are attached to the MurNAc residues. In growing bacteria, the PG sacculus undergoes continuous remodeling by PG synthases and PG hydrolases (6, 7). One of the PG hydrolases, <u>m</u>urein <u>l</u>ytic <u>t</u>ransglycosylase A (MltA) of *Escherichia coli*, is the focus of this report.

Lytic transglycosylases (LTs) cleave the β-1,4 MurNAc-GlcNAc glycosidic bond by an internal cyclization reaction that generates a cyclic 1,6-anhydroMurNAc end (anhMurNAc) (8). There are at least eight LTs in *E. coli* (for a recent review see (9)). Two of these enzymes have reasonably well-defined physiological roles. MltG is a terminase for PG glycan chain synthesis (10, 11), and Slt is uniquely important for preventing accumulation of uncrosslinked glycan strands generated during PG synthesis in the presence of β-lactam antibiotics (12). The precise physiological roles of the remaining LTs are less clear because overt morphological defects only become apparent when multiple genes are deleted (9, 13). This observation points to redundant and overlapping roles but does not resolve the specific contributions or relative importance of the various missing LTs in those mutants.

In *E. coli*, daughter cell separation is initiated by three *N*-acetylmuramoyl–L-alanine amidases that cleave the amide bond joining MurNAc to the first amino acid of the stem peptide, an L-alanine (14–16). Removal of peptide sidechains leaves behind peptide-free or “denuded” glycan strands (dnGs). The dnGs accumulate transiently in septal PG but are ultimately cleared by glycan hydrolases that have yet to be clearly defined in *E. coli* (17–19). Only three *E. coli* enzymes are known to turn over dnGs to any appreciable extent *in vitro*: the LTs MltA and MltE, and the muramidase/glucosaminidase DigH (20–22). Single gene deletion mutants of these enzymes divide normally, but more complex mutants lacking various combinations of these enzymes and other LTs exhibit a chaining phenotype indicative of delayed daughter cell separation (17, 23–25). Some of these mutants have been shown to accumulate dnGs (17, 25). Understanding who is responsible for turnover of septal dnGs is not simply a matter of enzymological bookkeeping. Rather, because dnGs serve as binding sites for division proteins with a SPOR domain, they play complex roles in coordinating multiple aspects of the constriction process (17, 26–32).

Here we report that MltA of *E. coli* contributes to turnover of septal dnGs. MltA is an outer membrane lipoprotein (20, 33). It is among the most abundant LTs in *E. coli* (34) and the most active *in vitro* when whole *E. coli* PG sacculi were used as substrate (35). Like most other LTs in *E. coli*, MltA is an exo-enzyme that releases GlcNAc-anhMurNAc disaccharides from the anhMurNAc end of PG glycans. It is active on PG glycans that have stem peptides, including crosslinked peptides, but also cleaves poly GlcNAc-MurNAc (i.e., dnGs) (20, 35). The biochemical attributes of MltA make it well-suited for clearing dnGs and other PG glycans from constriction sites. However, *in vivo* evidence for this function is limited. Although MltA is one of the missing LTs in some *E. coli* chaining mutants with multiple LT gene deletions (17, 23, 25), neither an *mltA* single mutant nor an *mltA mltB slt* triple mutant has a chaining phenotype (33).

## RESULTS

### MltA localizes to the *E. coli* division site, albeit weakly

We reasoned that if MltA processes septal PG it is likely to localize to the division site, so we used lambda Red recombination to fuse a HaloTag to the 3’ end of *mltA* at its native chromosomal locus (36, 37). The fusion strain and a wild-type control strain that lacks HaloTag were grown at 37°C in M9-glucose, then treated with JF549 dye to label the HaloTag. Cell extracts were analyzed by SDS-PAGE, which revealed a single fluorescent band at the expected size of ∼70 kDa, indicating the MltA-Halo fusion protein was produced and was stable (Fig. 1A). No such band was observed in extracts of the wild-type control strain. Microscopy of live cells revealed that MltA-Halo localized to the division site in about 40% of the cells (Fig. 1B). As expected, cells of the wild-type control strain were not fluorescent.

**Figure 1.**
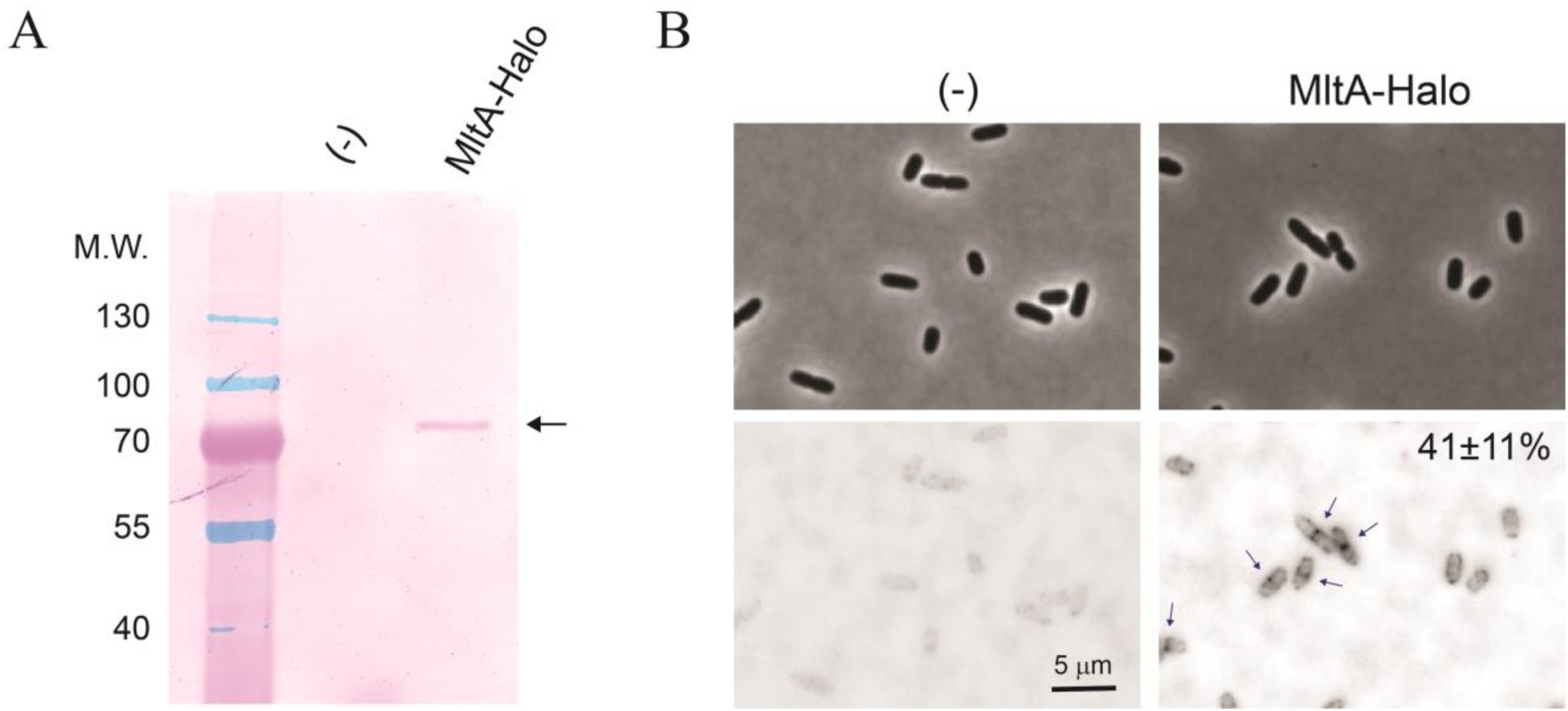
Septal localization of an MltA-Halo fusion protein produced from *mltA’s* native chromosomal locus. Cells of wild-type BW25113 (-) and an *mltA-halo* strain (EC5623) were grown in M9-glucose and treated with JF549 to label the Halo tag. (A) Proteins in whole cell extracts were separated by SDS-PAGE and the gel was imaged with a fluorescence scanner. Arrow points to the MltA-Halo fusion protein (predicted MW 75 kDa). Image is representative of five biological replicates. (B) Live cells were immobilized on agarose pads and imaged by phase contrast (above) and fluorescent (below) microscopy. Fluorescence micrographs were inverted to better visualize MltA-Halo. Arrows point to examples of septal localization. Numbers indicate the percentage of cells scored positive for septal localization (mean ± SD from three biological replicates).

The HaloTag proved somewhat cumbersome to work with because derivatization with JF549 requires several manipulations. We therefore fused MltA to two intrinsically fluorescent proteins, mCherry (mCh) and mNeonGreen (mNG). These fusions were expressed from a medium copy-number plasmid under control of a weak isopropyl-β-D-thiogalactopyranoside (IPTG) promoter. After testing several induction conditions, we settled on 20 μM IPTG, which resulted in ∼2.5-fold overproduction relative to the native MltA protein as assessed by Western blotting with antiserum against MltA (Fig. 2A, B). Fluorescence microscopy of live cells was used to assess septal localization of the fusions. Fluorescent signals were faint, so we inverted the gray scale in our micrographs to make localization sites easier to see. When cells were grown in LB, the MltA-mCh fusion localized to the midcell in ∼40% of cells. Localization increased to ∼50% of the cells when they were grown in LB0N, which is LB made without NaCl (Fig. 2C). The MltA-mNG fusion also localized to the midcell in about half of the population, although we did not quantify this carefully (Fig. 2D). Substituting MltA’s catalytic aspartate with alanine had little or no effect on localization (Fig. 2C) (38).

**Figure 2.**
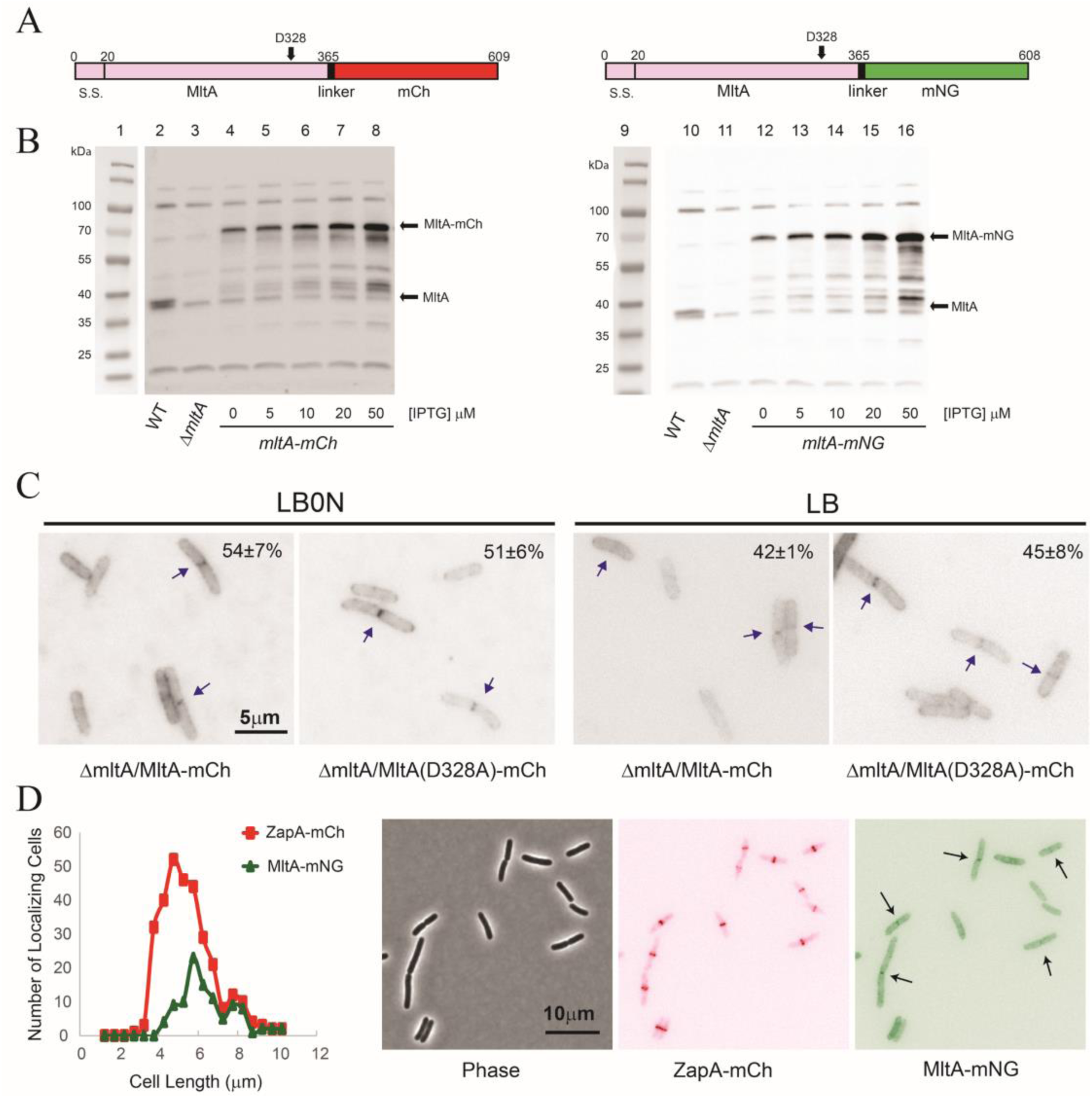
Septal localization of fluorescent MltA fusion proteins produced from plasmids. (A) Schematic of the MltA-mCh and MltB-mNG fusion proteins. S.S., signal peptide with Signal Peptidase II cleavage site following residue 20, an alanine. D328 is an essential catalytic aspartate. (B) Western blot analysis of MltA-mCh and MltA-mNG stability and abundance. Cells were grown in LB0N containing IPTG as indicated to induce expression of the fusions. Proteins were subjected to SDS-PAGE and Western blot with polyclonal anti-MltA serum. Native MltA runs slightly above a cross-reactive band of unknown origin also present in the Δ*mltA* sample. Molecular mass standards are on the left. WT, EC251; Δ*mltA*, EC3546; *mltA-mCh*, EC6334; *mltA-mNG*, EC5801. (C) Fluorescence micrographs of cells producing either wild-type or catalytically inactive MltA-mCh fusion proteins. Arrows point to examples of septal localization. Numbers in the upper right indicate the percentage of cells scored positive for septal localization (mean ± SD from three biological replicates). Cells were grown in LB0N or LB as indicated containing 20 μM IPTG. (D) Cells of strain EC5868 were measured and scored for the presence of a red (ZapA-mCh) or green (MltA-mNG) band at the midcell. Data were pooled from two experiments with >300 cells scored per experiment. Representative phase contrast and fluorescence micrographs are shown on the right with arrows indicating examples of MltA-mNG localization.

### MltA is a late recruit to the division site

In *E. coli*, cell division is mediated by a collection of over 30 proteins that accumulate into a loosely defined structure commonly referred to as the divisome or septal ring (39–41). Divisome assembly begins with FtsZ and its associated proteins like FtsA, ZipA and ZapA. Proteins more directly involved in PG synthesis such as FtsQ, FtsW, FtsI and FtsN are recruited later (42). Visible constriction coincides with the arrival of these late proteins, especially FtsN, a key activator of septal PG synthesis (28, 32, 43–46). Our observation that MltA localizes in only 40-50% of the cells suggested it is a late recruit, which is compatible with a role in removing dnGs.

To compare MltA localization to that of an early recruit, we transformed our *mltA-mNG* plasmid into a strain that expresses *zapA-mCherry* (ZapA-mCh) from *zapA’s* native locus on the chromosome. Using cell length as a proxy for age, ZapA-mCh localized well ahead of MltA-mNG (Fig. 2D). We then asked whether septal localization of MltA requires the presence and/or function of FtsZ, FtsQ, FtsI and FtsN. These experiments were done by introducing the *mltA-mCh* plasmid into temperature-sensitive (Ts) or depletion strains of the Fts proteins. In all cases, MltA-mCh localization was lost under non-permissive conditions (Fig. 3). Thus, by two criteria, timing and dependency, MltA is a late recruit to the division site.

**Figure 3.**
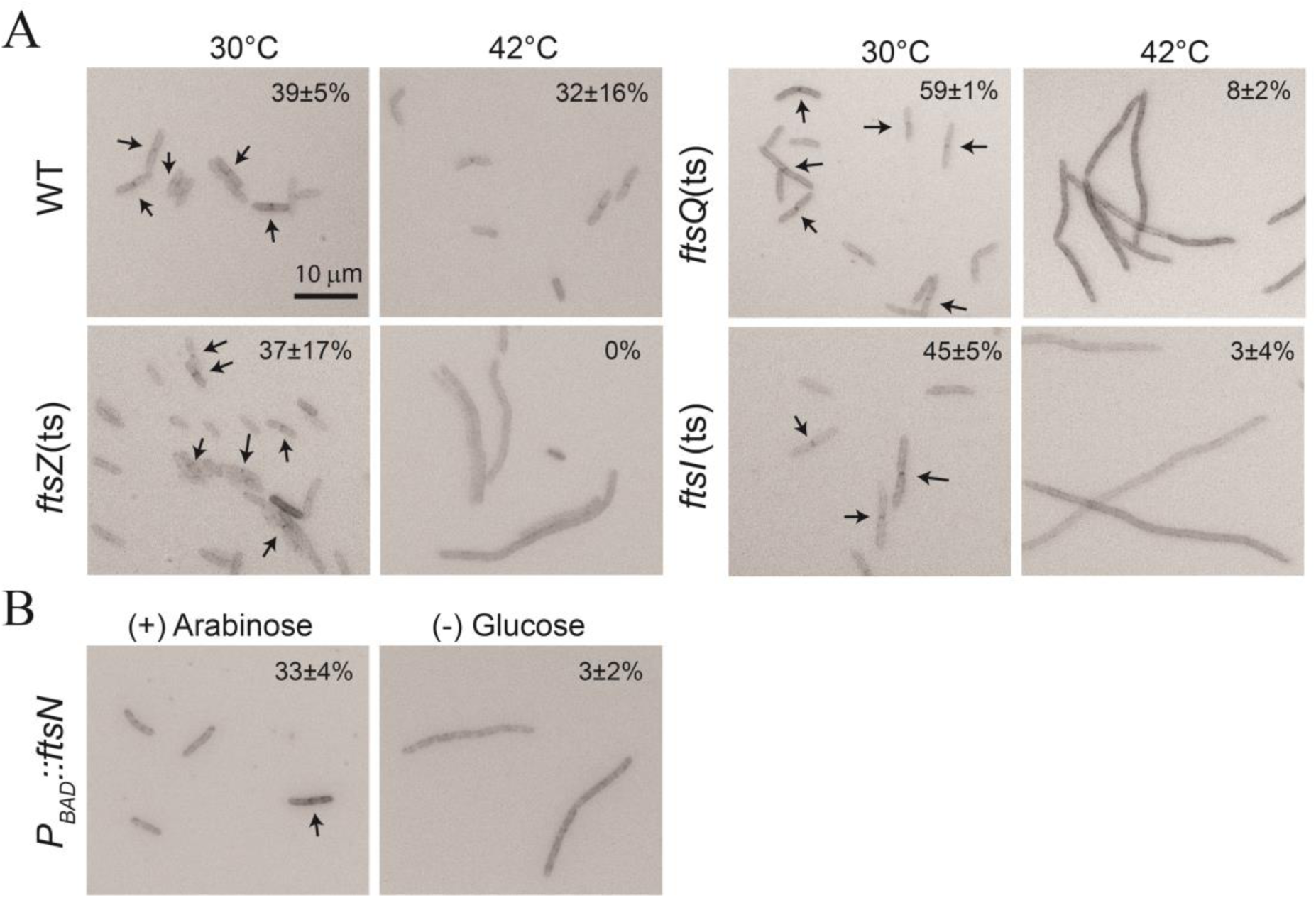
Dependency of MltA-mCh localization on other division proteins. Representative fluorescence micrographs of wild-type and *fts* mutant strains producing MltA-mCh from a plasmid. (A) To test Ts mutants, strains grown at 30°C to OD_600_ ∼0.5 were back diluted 1:10 and grown for 60-90 minutes at either 30°C or 42°C. Live cells were immobilized on agarose pads for microscopy. (B) For the FtsN-depletion stain, the growth media contained 0.5% arabinose (*ftsN* expressed) or 0.2% glucose (*ftsN* not expressed). Percentages are mean ± SD of cells scored positive for septal localization from two experiments. Arrows indicate examples of localization. Strains shown are WT, EC6447; *ftsZ*(ts), EC6449; *ftsQ*(ts), EC6366; EC6362, *ftsI*(ts) and EC6681, *P_BAD_::ftsN*.

### MltA contributes to turn over of denuded glycans in septal PG *in vivo*

If MltA has a major role in promoting daughter cell separation, a Δ*mltA* mutant should have a chaining phenotype. However, an *mltA* mutant was said to have normal morphology (33), so if there are any defects in daughter cell separation they are likely to be subtle. We therefore grew a Δ*mltA* mutant in a low osmotic strength media (LB0N) known to exacerbate defects in LT mutants (18, 19, 47, 48). However, the Δ*mltA* mutant grew at the same rate as WT and was not elongated or chained even after 12 mass doublings to allow time for subtle defects to become manifest (Fig. 4A,B). In fact, the mutant was ∼10% shorter than wild-type (5.0 ± 0.1 µm versus 5.6 ± 0.1 µm, p < 0.05, n =3 experiments), and staining membranes with FM4-64 revealed that fewer cells were in the process of dividing (34 ± 6% versus 41 ± 2%, p < 0.05, n = 3 experiments). Therefore, based on cell morphology, loss of MltA slightly impairs elongation but not daughter cell separation.

**Figure 4.**
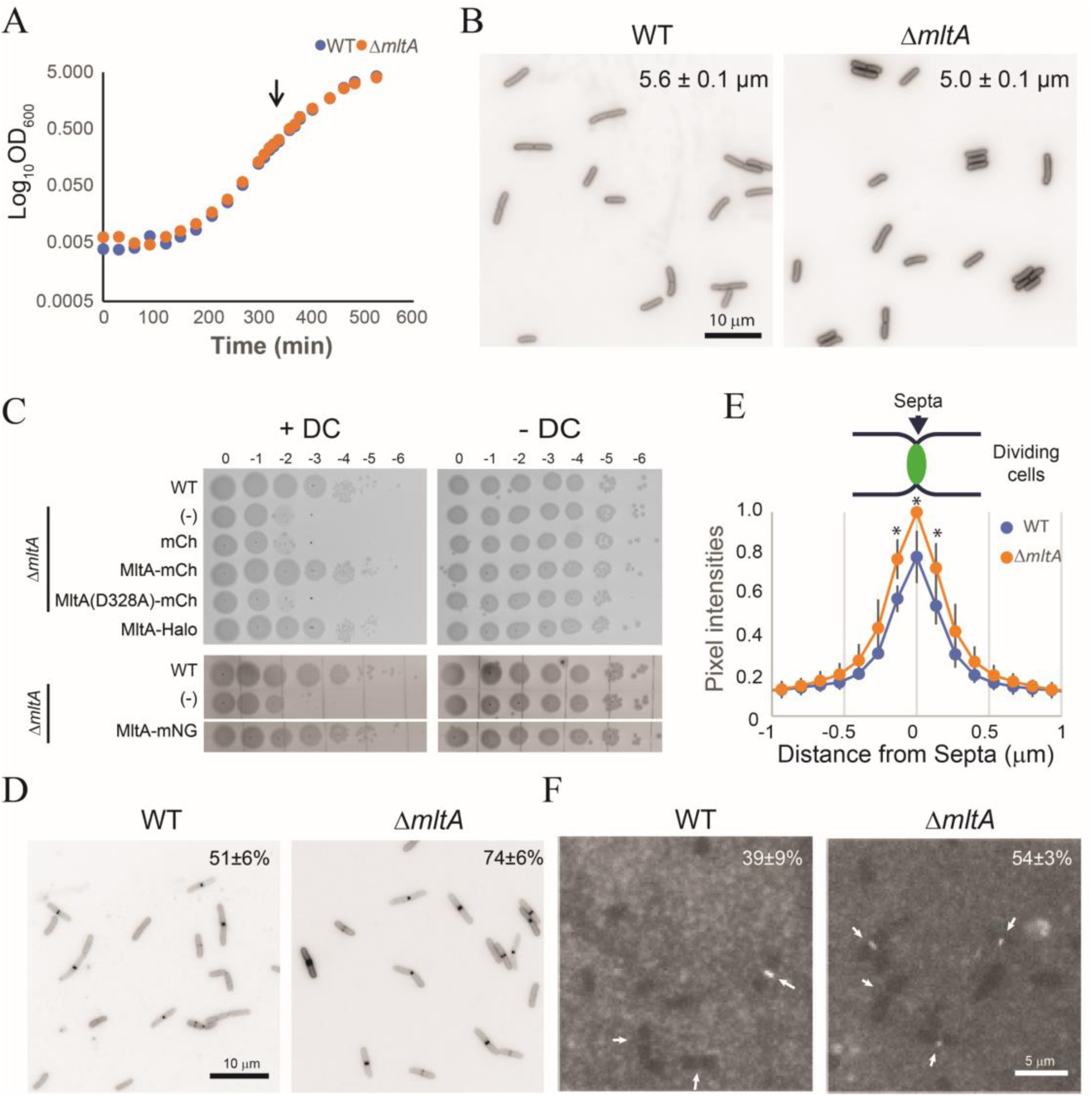
Phenotypic characterization of a Δ*mltA* strain. (A) Growth of WT (EC251) and Δ*mltA* (EC3546) in LB0N. (B) At OD_600_ = 0.5 (arrow) samples were fixed, membranes were stained with FM4-64, and cells were photographed with fluorescence microscopy. Cell lengths in the upper right of each panel are the mean ± SD from three experiments with >100 cells measured per experiment. (C) Deoxycholate resistance evaluated by spot titer assay. Ten-fold serial dilutions of the indicated strains were spotted onto LB0N plates containing 20 µM IPTG with or without 0.1% deoxycholate. Plates were photographed after 18 h at 37°C. The lower set of strains were on the same plate, but the photograph was cropped to remove non-relevant strains. Strains are WT, EC251: (-), EC3546; mCh, EC6475; MltA-mCh, EC6334; MltA(D328A)-mCh, EC6332; MltA-Halo, EC5623; MltA-mNG, EC5801. (D) Fluorescence micrographs of WT (EC6534) or Δ*mltA* (EC6539) producing ^TT^GFP-FtsN^SPOR^. The percentage of cells scored positive for septal localization is the mean ± SD from 3 experiments with >100 cells scored per experiment. (E) Abundance of septal ^TT^GFP-DamX^SPOR^. Data are graphed as the mean ± SD based on transects from at least 45 cells scored positive for septal localization. *p < 0.05 in a Student’s t-test. (F) Fluorescence micrographs of GFP-DamX^SPOR^ bound to PG sacculi from WT (EC251) or Δ*mltA* (EC3546). Purified PG sacculi were immobilized on a glass slide, incubated with 10 nM GFP-DamX^SPOR^ protein for 30 min, washed and imaged by fluorescence microscopy. Percentages indicate the fraction of sacculi scored positive for septal localization (mean ± SD from 2 experiments with >100 cells score per experiment).

Another phenotype associated with inefficient turnover of septal PG is deoxycholate sensitivity (23), presumably because residual septal PG interferes with constriction of the outer membrane, which disrupts its important barrier function (49). MltA was previously found to make a small contribution to deoxycholate resistance in a high-throughput phenotypic analysis of the Keio collection (50). We confirmed this result in a different assay format using spot titers on LB0N plates containing 0.1% deoxycholate (Fig. 4C). We then took advantage of the deoxycholate sensitivity phenotype to test the functionality of our various fluorescent and HaloTag fusions, all of which rescued viability, indicating they are functional (Fig. 4C). However, substitution of the catalytic aspartate with alanine (D328A) abrogated rescue, as expected (Fig. 4C).

While deoxycholate sensitivity is suggestive of a division defect, it is not definitive because defects anywhere in the outer membrane can allow deoxycholate to enter the periplasm. To ask directly whether dnGs accumulate in the Δ*mltA* mutant, we assayed septal localization of a green fluorescent protein (GFP) fusion to the SPOR domain from FtsN. SPOR domains bind dnGs, so their abundance is proportional to the amount of GFP-FtsN^SPOR^ fluorescence at the septum (17, 26, 28, 30, 32). We observed that the fraction of cells scored positive for septal localization of GFP-FtsN^SPOR^ increased from about 50% in a WT background to about 75% in the Δ*mltA* mutant (Fig. 4D). We also observed an increase of ∼20% in the intensity of septal fluorescence in cells scored positive for septal localization (i.e., omitting from this measurement cells without a fluorescent band at the midcell) (Fig. 4E). This finding was replicated in an in vitro assay in which we observed increased binding of GFP-DamX(SPOR) protein to septal regions of PG sacculi isolated from a Δ*mltA* mutant as compared to WT (Fig. 4F). We conclude that MltA makes a small but measurable contribution to removing dnGs from septal PG.

### MltA binds to septal regions of purified *E. coli* PG sacculi

To visualize binding of MltA to PG sacculi *in vitro*, we purified a GFP-MltA fusion protein with a D328A substitution to prevent the PG from being digested during the assay. For comparison purposes, we isolated sacculi from three *E. coli* strains: wild-type, Δ*mltA*, and Δ*LTs*. In the latter, septal dnG is at least 20-fold more abundant than in wild-type owing to the deletion of genes for five known and one potential LT (Δ*mltACDE* Δ*slt* Δ*rlpA*) (17). Representative fluorescence micrographs are shown in Figure 5A and an additional example of binding to Δ*LTs* sacculi is shown in Figure 5B. The sacculi appeared dark against a lighter background. It was very difficult to visualize GFP-MltA(D328A), which appeared as a faint whitish band of fluorescence at some constriction sites. The fraction of sacculi exhibiting septal localization was 33% for wild-type (162 sacculi) and to 48% for Δ*mltA* (148 sacculi). The Δ*LTGs* mutant was not be scored because we could not reliably distinguish bound fusion protein from deep constrictions. In any case, fluorescence was not noticeably brighter with sacculi from the Δ*LTs* strain. These findings indicate MltA binds preferentially to septal PG, this binding is weak and/or there are very few binding sites for MltA in septal PG, and the PG structure to which MltA binds is not simply dnGs because if it were, fluorescence signals would have been more prominent with sacculi from the Δ*LTs* strain.

**Figure 5.**
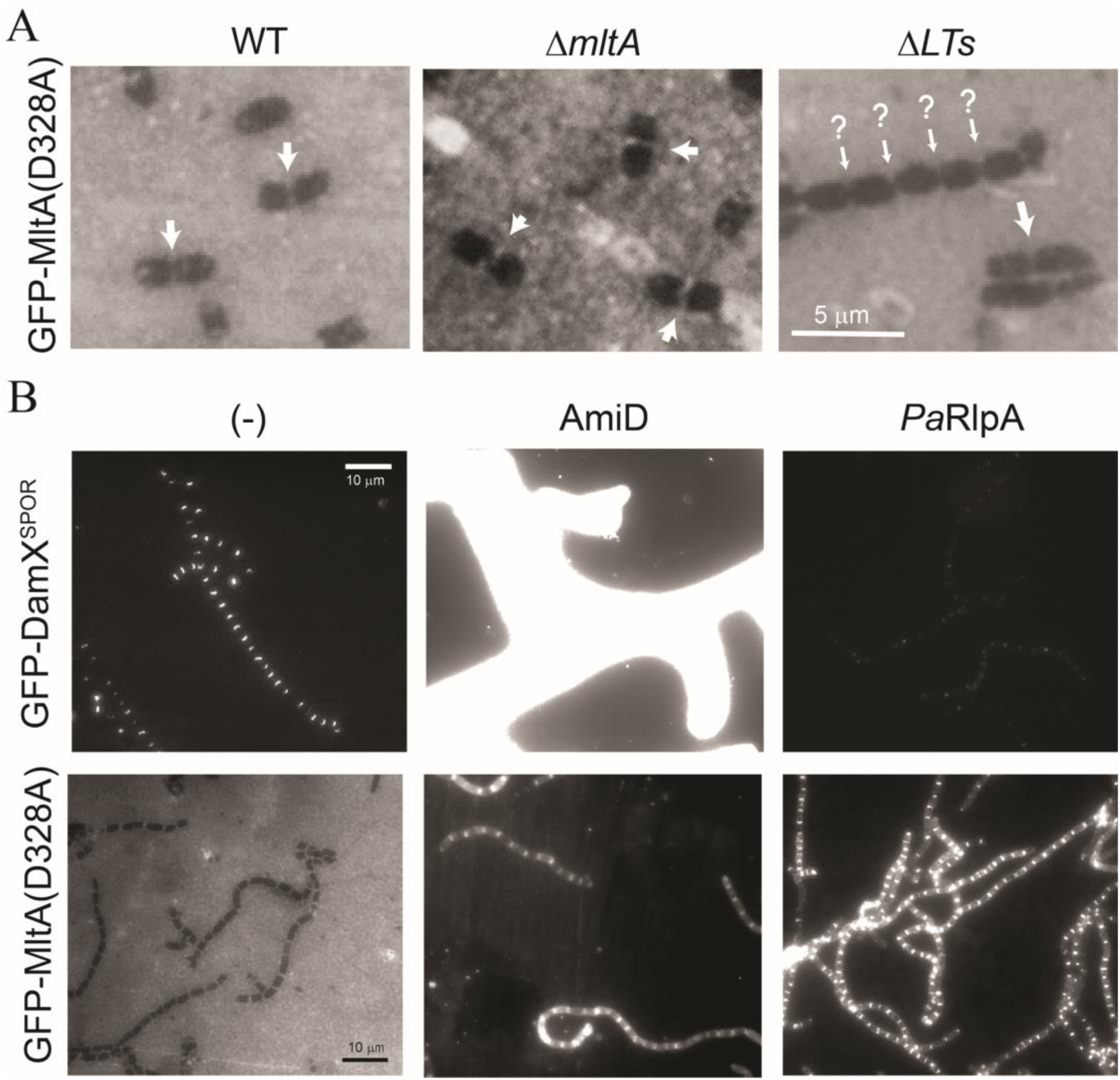
An MltA-GFP fusion protein binds to septal regions of PG sacculi purified from *E. coli*. (A) Representative fluorescence micrographs of MltA(D328A)-GFP bound to immobilized PG sacculi from wild-type and mutant *E. coli* strains as indicated. MltA(D328A)-GFP was added at 100 nM, incubated for 30 min, and sacculi were washed before imaging. Arrows indicate weak GFP signals; strong signals were never observed. Question marks indicate some evidence for binding but this is uncertain. (B) MltA may bind to anhMurNAc ends of dnGs. Immobilized sacculi from the Δ*LTs* strain were treated for 1 h with 2.5 µM AmiD or 5 µM RlpA from *P. aeruginosa*. AmiD is a cell wall amidase that removes stem peptides. *Pa*RlpA is an endo-LT that removes dnGs while leaving behind glycan strands with anhMurNAc ends. PG digestion was terminated by washing the sacculi, which were then tested as binding substrates for 100 nM MltA(D328A)-GFP or 10 nM GFP-DamX^SPOR^. Assays included a no-enzyme control (-). Micrographs are representative of three experiments. Sacculi were purified from WT, EC251; Δ*mltA*, EC3546; and Δ*LTs*, EC3708.

### Evidence that MltA binds preferentially to 1,6-anhMurNAc ends of glycan chains

Although we were skeptical that MltA localizes to septal PG by binding to dnGs, we performed some additional experiments to test this possibility. If a dnG is MltA’s preferred binding site, removing stem peptides by treating sacculi with AmiD should increase binding, while removing dnGs by treating sacculi with RlpA from *Pseudomonas aeruginosa* (*Pa*RlpA) should decrease binding. Indeed, this is what we observed with the control protein GFP-DamX^SPOR^, which binds to dnGs (Fig. 5B, note all micrographs adjusted identically for direct comparisons) (17). However, GFP-MltA(D328A) behaved differently. Limited digestion with AmiD resulted in only a small increase in binding, while limited digestion with *PaRlpA* caused a much larger increase, especially in septal regions (Fig. 5B). What could *Pa*RlpA be doing that would increase the number of binding sites for MltA? *PaRlpA* is a dnG-specific endo-LT, so in addition to removing dnGs, it creates anhMurNAc ends. Thus, the simplest explanation for our observations is that MltA binds to the anhMurNAc ends of PG glycans, which makes sense biochemically because MltA is an exo-LT that removes disaccharides from the anhMurNAc end of PG glycans (20).

## DISCUSSION

LTs are important for a variety of processes in bacteria, including elongation, division, making holes in the PG for trans-envelope complexes, PG recycling and preventing PG turnover products from accumulating to deleterious levels in the periplasm (for reviews see (9, 13, 51, 52). However, linking specific LTs to any of these processes can be difficult because functional redundancy obscures phenotypic defects in single gene deletion mutants.

Here we determined that MltA contributes to removal of dnGs from septal PG during cell division in *E. coli*. Prior studies of MltA had suggested this might be the case based on the findings that MltA has appropriate substrate specificity *in vitro* and mutants lacking multiple LTs (including MltA) exhibit chaining phenotypes and accumulate dnGs in septal PG (17, 20, 23, 25). But loss of MltA alone is not sufficient to cause a chaining defect, and even an *mltA mltB slt* triple mutant was found to have normal morphology (33). We made two observations that establish MltA as a *bona fide* septal LT. First, fusing HaloTag to the 3’ end of *mltA* at its native chromosomal locus revealed weak but convincing septal localization (Fig. 1). Second, there is a small but reproducible accumulation of dnGs in an *mltA* deletion mutant even though all other LTs are present (Fig. 4).

We do not think that MltA is restricted to processing septal PG, and the fact that a simple Δ*mltA* mutant does not grow as chains of linked daughter cells indicates additional LTs turn over septal PG in *E. coli*. We suggest that future studies should prioritize MltE, DigH and MltC. MltE and DigH utilize dnG substrates, but MltC does not (21, 22, 53). However, *mltCE* double mutants of *Salmonella* Typhimurium and *E. coli* have chaining phenotypes under some growth conditions (24, 47). Finally, there is the enigmatic RlpA protein, which is clearly a dnG-specific LT important for daughter cell separation in *Pseudomonas aeruginosa* and *Vibrio cholera* (18, 19). But *E. coli* RlpA has a degenerate active site and experimental evidence indicates it has probably lost enzymatic function (18).

We found that MltA is a late recruit to the divisome. Whether MltA is recruited by binding a divisome protein or a PG structure remains to be determined. Our *in vitro* binding assays favor the latter possibility and suggest this structure could be the anhMurNAc ends of PG glycans, which are created by LTs. But for anhMurNAc to drive recruitment, these structures would have to be enriched in septal PG. It’s not obvious why that would be the case. Another interesting question concerns how the activity of MltA is regulated. Given MltA’s broad substrate specificity, it ought to breakdown PG glycans everywhere in the cell and at all times, which would be lethal. The fact that MltA is tethered to the outer membrane probably limits its access to the sacculus, so one level of control might be mediated via constriction of the outer membrane. But additional regulation is presumably required to explain how MltA activity is controlled in time and space.

## MATERIALS AND METHODS

### Media

Lysogeny broth (LB) contained per liter 10 g tryptone, 5 g yeast extract, 10 g NaCl and 15 g agar (for plates). NaCl was omitted for LB0N. M9-glucose minimal medium contained 0.4% D-glucose, 1x MEM amino acids and 1x MEM vitamins as described previously (45). Antibiotics were used at the following concentrations: ampicillin (Amp), 200 µg/ml; kanamycin (Kan), 40 µg/ml; spectinomycin (Spc), 100 μg/ml. Isopropyl-β-D-1-thiogalactopyranoside (IPTG) was used at 20 µM to induce genes under control of modified *Trc* promoters unless indicated otherwise. L-Arabinose (Ara) and D-glucose (Glu) were used at 0.2% to regulate expression of genes under *P_BAD_* control.

### Bacterial strains, plasmids and oligonucleotides

Strains, plasmids and oligonucleotides are listed in Tables 1-3. Strain construction followed published procedures for P1 transduction, Lambda Red recombineering and transformation (36, 54). All strains are listed in Table 1. Strains new to this study were constructed as follows. *EC3496*: P1 transduction from JW2784-1 into EC251, selecting Kan^r^. *EC3546*: Used pCP20 to evict the Kan^r^ marker from EC3496. EC5546: Constructed by λ Red recombination. Primers P2591 and P2592 were used to PCR-amplify a 2.3 kb DNA fragment from template plasmid pDSW2161. The PCR product encoded the 3’ end of *mltA* fused to HaloTag, a Kan^r^ cassette and downstream homology for recombination at the native *mltA* locus. This DNA fragment was introduced into BW25113/pKD46 by electroporation, selecting Kan^r^. *EC5594*: Constructed by λ Red recombination. Primers P2627 and P2628 were used to amplify a 2.2 kb DNA fragment from template plasmid 2179. The PCR product encoded the 3’ end of *zapA* fused to *mCherry*, a Kan^r^ cassette, and downstream homology for recombination at the native *zapA* locus. This DNA fragment was introduced into BW25113/pKD46 by electroporation, selecting Kan^r^. *EC5623*: P1 transduction from EC5546 into EC251, selecting Kan^r^. All other strains were constructed by transformation of the indicated plasmid into the indicated strain background.

**TABLE 1.**
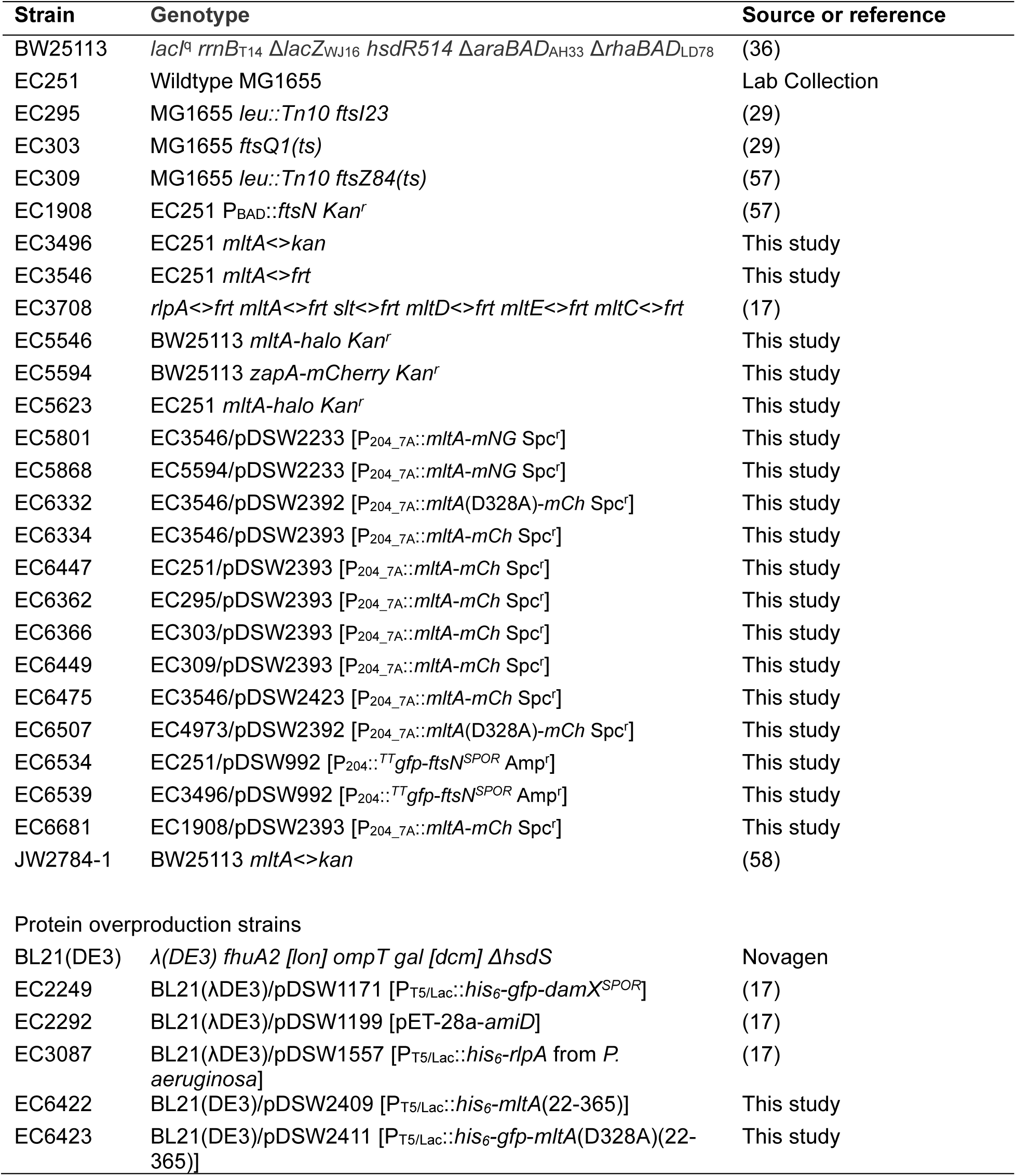
Strains used in this study.

**TABLE 2.** Plasmids used in this study.

| Plasmid | Relevant features | Source or reference |
| --- | --- | --- |
| pACYC184 | p15A Cam<sup>r</sup> Tet<sup>r</sup> | (59) |
| pCP20 | λP<sub>R</sub>::*FLP* λcl857 *bla cat* Rep<sup>TS</sup> (pSC101 derivative) | (60) |
| pDSW204 | IPTG inducible (P<sub>204</sub>); *lacI<sup>q</sup>* pBR ori Amp<sup>r</sup> | (61) |
| pDSW912 | *rfp* fusion vector (P<sub>204</sub>::MCS-*mCherry*) *lacI<sup>q</sup>* pBR ori Amp<sup>r</sup> | (29) |
| pDSW913 | *rfp* fusion vector (P<sub>206</sub>::MCS-*mCherry*) *lacI<sup>q</sup>* pBR ori Amp<sup>r</sup> | (29) |
| pDSW992 | P<sub>204</sub>::*TTgfp-ftsN<sup>SPOR</sup>*(240–319) *lacI<sup>q</sup>* pBR ori Amp<sup>r</sup> | (62) |
| pDSW1171 | P<sub>T5/Lac</sub>::*his6-gfp-damX<sup>SPOR</sup>*(338–428) *lacI<sup>q</sup>* pBR ori Amp<sup>r</sup> | (17) |
| pDSW1557 | P<sub>T5/Lac</sub>::*his6-ParlpA* (28–341) *lacI<sup>q</sup>* pBR ori Amp<sup>r</sup> | (17) |
| pDSW1642 | P<sub>206</sub>::*drpB-gfp* *lacI<sup>q</sup>* pBR ori Amp<sup>r</sup> | (63) |
| pDSW1877 | P<sub>204_7A</sub>::*gfp-MCS* *lacI<sup>q</sup>* attP<sub>Φ80Y</sub> oriR<sub>R6KY</sub> Spc<sup>r</sup> | (55) |
| pDSW1926 | P<sub>204_7A</sub>::*mNG-ftsN*, *lacI<sup>q</sup>* attP<sub>Φ80Y</sub> oriR<sub>R6KY</sub> Spc<sup>r</sup> | (45) |
| pDSW1959 | P<sub>206</sub>::*drpB-gfp frt-Kan-frt* *lacI<sup>q</sup>* pBR ori Amp<sup>r</sup> Kan<sup>r</sup> | This study |
| pDSW2132 | P<sub>204_7A</sub>::*SSdsbA-cys-(KpnI site)-Halo* *lacI<sup>q</sup>* pSC101 ori Kan<sup>r</sup> | This study |
| pDSW2142 | P<sub>204_7A</sub>::*SSdsbA-mltA-Halo* *lacI<sup>q</sup>* pSC101 ori Kan<sup>r</sup> | This study |
| pDSW2161 | P<sub>206</sub>::*Halo frt-Kan-frt* *lacI<sup>q</sup>* pBR ori Amp<sup>r</sup> Kan<sup>r</sup> | This study |
| pDSW2179 | P<sub>206</sub>::*mCherry frt-Kan-frt* *lacI<sup>q</sup>* pBR ori Amp<sup>r</sup> Kan<sup>r</sup> | This study |
| pDSW2229 | P<sub>204_7A</sub>::*gfp* *lacI<sup>q</sup>* p15A ori oriR<sub>R6KY</sub> Spc<sup>r</sup> | This study |
| pDSW2233 | P<sub>204_7A</sub>::*mltA-mNeonGreen* p15A ori oriR<sub>R6KY</sub> Spc<sup>r</sup> | This study |
| pDSW2235 | P<sub>204_7A</sub>::*mltA-mCherry* p15A ori oriR<sub>R6KY</sub> Spc<sup>r</sup> | This study |
| pDSW2240 | P<sub>204_7A</sub>::*mltA<sup>D328A</sup>-mNeonGreen* p15A ori oriR<sub>R6KY</sub> Spc<sup>r</sup> | This study |
| pDSW2392 | P<sub>204_7A</sub>::*mltA<sup>D328A</sup>-mCherry* p15A ori oriR<sub>R6KY</sub> Spc<sup>r</sup> | This study |
| pDSW2393 | P<sub>204_7A</sub>::*mltA-mCherry* p15A ori oriR<sub>R6KY</sub> Spc<sup>r</sup> | This study |
| pDSW2411 | P<sub>T5</sub>::*His6-gfp-mltA<sup>D328A</sup>*(22–365) *lacI<sup>q</sup>* pBR ori Amp<sup>r</sup> | This study |
| pDSW2423 | P<sub>204_7A</sub>::*mCh* p15A ori oriR<sub>R6KY</sub> Spc<sup>r</sup> | This study |
| pET-28a-amiD | P<sub>T7</sub>::*his6-amiD* *lacI<sup>q</sup>* pBR ori Kan<sup>r</sup> | (64) |
| PJL74 | P<sub>204_7A</sub>::*SSdsbA-Halo-ftsN*(243–319) *lacI<sup>q</sup>* oriR<sub>R6KY</sub> attP<sub>Φ80</sub> Spc<sup>r</sup> | (45) |
| pKD13 | oriR<sub>R6KY</sub> *frt-kan-frt* Amp<sup>r</sup> | (36) |
| pKD46 | P<sub>BAD</sub>::*gam beta exo araC* pSC101 ori(Ts) Amp<sup>r</sup> | (36) |
| pTH18-kr | IPTG-inducible (P<sub>Iac</sub>), Kan<sup>r</sup>, pSC101 *ori* | (65) |

**TABLE 3.**
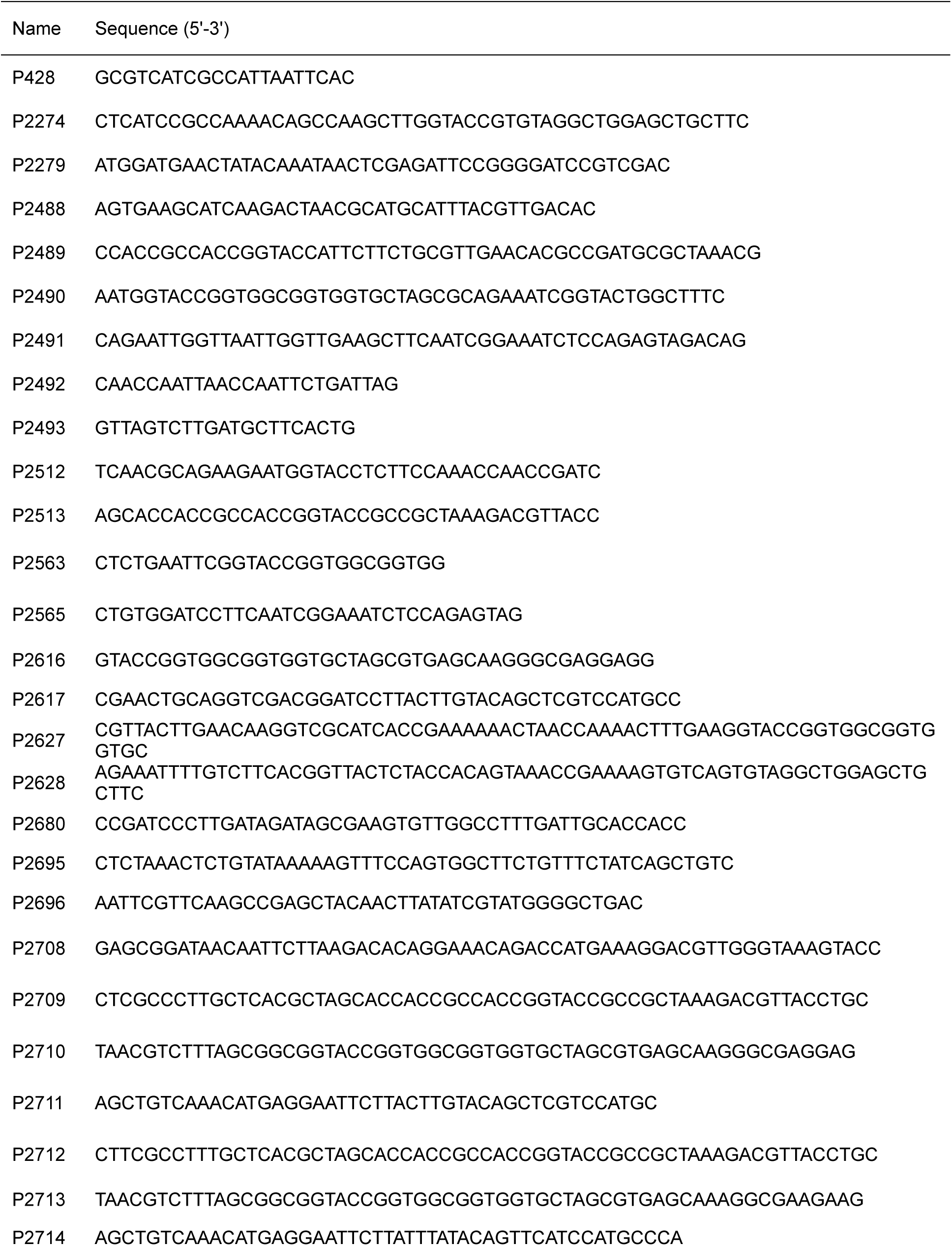
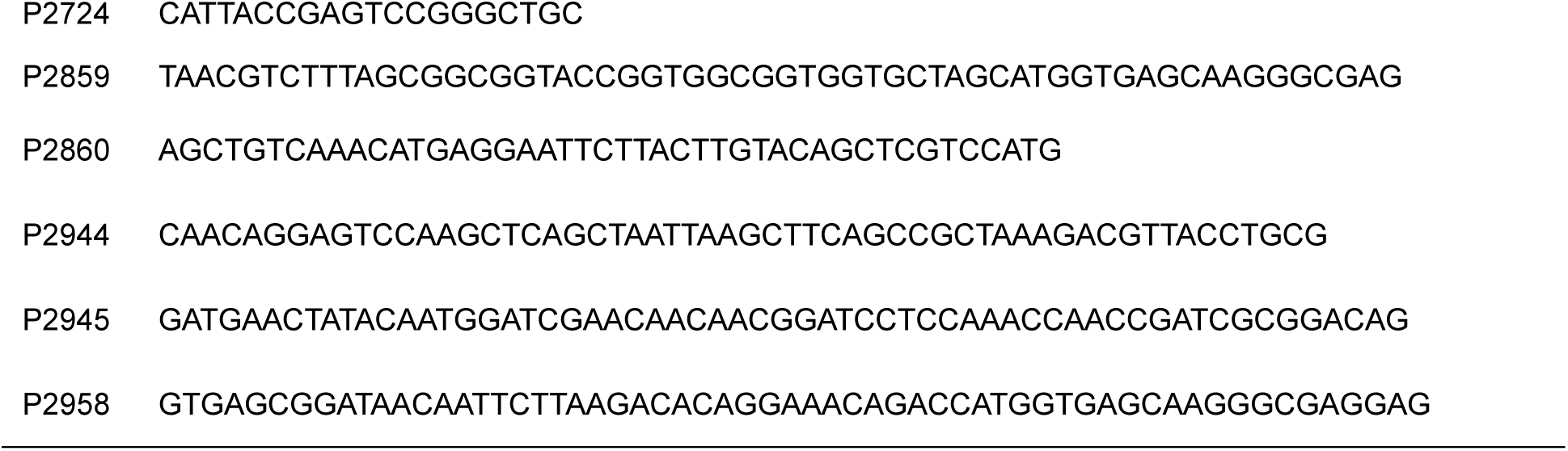
Oligonucleotides used in this study.

Plasmid construction used restriction enzymes, T4 DNA ligase, Q5 DNA polymerase, and NEBuilder HiFi DNA Assembly Master Mix from New England Biolabs (NEB, Ipswich, MA). Plasmid DNA was purified using the QIAprep Spin Miniprep Kit (Qiagen, Germantown, MA). Monarch kits from NEB were used to recover DNA fragments from PCR reaction mixtures and agarose gels. Regions of plasmids constructed using PCR were verified by dye-termination cycle-sequencing performed by the DNA Core Facility of the Carver College of Medicine. Details for construction of specific plasmids are provided in the following paragraph.

*pDSW1959*: PCR amplified *frt-kan-frt* from pKD13 with primers P2279 and P2274. The 1362 bp fragment was inserted by isothermal assembly into HindIII-digested pDSW1642. *pDSW2423*: PCR amplified *mCherry* from pDSW2235 with primers P428 and P2958. The 805 bp product was inserted by isothermal assembly into EcoRI/AflII-digested pDSW2235. *pDSW2132*: Isothermal assembly of three DNA fragments generated as follows: (i) A 1600 bp PCR product including *lacI* and the Tat system signal sequence *dsbA* was amplified from pJL74 with primers P2488 and P2489. (ii) A 943 bp PCR product with HaloTag was amplified from pJL74 with primers P2490 and P2491. (iii) A 2522 bp PCR product including the backbone of pTH18Kr was amplified from pTH18Kr with primers P2492 and P2493. *pDSW2142*: PCR amplified *mltA* from EC251 genomic DNA with primers P2512 and P2513. The 1074 bp product was inserted by isothermal assembly into KpnI-digested pDSW2132. *pDSW2161*: PCR amplified HaloTag from pDSW2142 with primers P2563 and P2565. The 936 bp was digested with EcoRI and BamHI, then ligated into the same sites of pDSW1959. *pDSW2179*: PCR amplified *mCherry* from pDSW912 with primers P2616 and P2617. The 753 bp PCR product inserted by isothermal assembly into NheI/BamHI-digested pDSW2161. *pDSW2229*: PCR amplified the p15A origin from pACYC184 with primers P2695 and P2696. The 1042 bp PCR product was inserted by isothermal assembly into NheI/NotI-digested pDSW1877. *pDSW2233*: Isothermal assembly of three DNA fragments generated as follows. (i) A 1172 bp PCR product including *mltA* was amplified from EC251 genomic DNA with primers P2708 and P2712. (ii) A 770 bp PCR product encoding *mNeonGreen* was amplified from pDSW1926 with primers P2713 and P2714. (iii) pDSW2229 was digested with AflII/EcoRI. *pDSW2235*: Three fragment isothermal assembly. (i) A 1171 bp PCR product encoding *mltA* was amplified from EC251 genomic DNA with primers P2708 and P2709. (ii) A 770 bp PCR product encoding *mCherry* was amplified from pDSW913 with primers P2710 and P2711. (iii) pDSW2229 digested with AflII/EcoRI. *pDSW2240*: PCR amplified ‘*lacI^q^*-*mltA(D328A)’* from pDSW2233 with primers P2724 and P2680. The 1626 bp product was inserted into EcoRV-digested pDSW2233. *pDSW2392*: PCR amplified *mCherry* from pDSW913 with primers P2859 and P2860. The 773 bp product was inserted by isothermal assembly into KpnI/EcoRI-digested pDSW2240. *pDSW2393*: PCR amplified *mCherry* from pDSW913 with primers P2859 and P2860. The 773 bp product was inserted by isothermal assembly into KpnI/EcoRI-digested pDSW2233. *pDSW2411*: PCR amplified *mltA*(D328A) from pDSW2240 with primers P2944 and P2945. The 1099 bp product was inserted by isothermal assembly into BamHI/EcoRI-digested pDSW2240.

### Experiments using MltA-Halo

Strains were grown overnight in LB at 37°C. Cells from overnight cultures were pelleted and resuspended in M9-glucose supplemented with amino acids and vitamins. Cultures were then diluted 1:2000 into 25 ml of the same media in a 250 ml Erlenmeyer baffled culture flask, and grown at 37°C, 210 rpm to OD_600_ ∼0.35. For detection of MltA-Halo by SDS-PAGE, a 100 µl aliquot of culture was incubated with 1 μM of JF549 HaloTag ligand and processed as described (55). The high concentration of JF549 improved signals in the gels. For microscopy, a 100 µl aliquot of the culture was incubated with 100 nM JF549. The low concentration of JF549 reduced background fluorescence and thus improved signals in microscopy. After 30 min incubation at room temperature in the dark, cells were washed with PBS and fixed with 2.5% paraformaldehyde and 0.8% glutaraldehyde buffered with NaPO_4_. Cells were washed to remove fixing reagents prior to mounting on agarose pads for microscopy (18).

### Western blotting

Starter cultures were grown overnight at 30°C in LB (EC251, EC3546) or LB-Spc (EC5801, EC6334). The next morning, cultures were diluted 1:200 in LB or LB-Spc containing various concentrations of IPTG as indicated in Figure 2B and grown at 30°C to OD_600_ ∼0.5. Cells from 1 ml of culture were harvested by centrifugation and the cell pellet was taken up in 100 µl of 1x Laemmli Sample Buffer. Samples were heated for 10 min at 95°C before loading 10 µl onto a precast mini-PROTEAN TGX gel (10% polyacrylamide, from Bio-Rad, Hercules, CA). Electrophoresis, transfer to nitrocellulose, and blot development followed standard procedures. Blocking was with 5% nonfat dry milk diluted in Tris-buffered saline containing 0.1% TWEEN 20 (TBST). Primary antibody was rabbit anti-MltA serum that had been incubated with an extract from a Δ*mltA* strain, EC3546, to reduce cross-reaction with cellular proteins (56). The preabsorbed antiserum was used at a dilution of 1:1000 in TBST incubated with the blot for two hours. Secondary antibody was horseradish peroxidase-conjugated goat anti-rabbit antibody diluted 1:8000 in TBST and used for two hours (Pierce, Rockford, IL). Detection was with SuperSignal Pico West chemiluminescent substrate (Pierce, Rockford, IL). Blots were visualized with ChemDoc Touch imaging system (BioRad, Hercules, CA)

### Growth of fluorescent protein fusion strains for microscopy

(i) MltA-mCh in Fig. 2C. EC6332 or EC6334 were grown overnight in LB-Spc with 10 μM IPTG at 30°C. The next day, cultures were diluted 1:200 into LB-Spc or LB0N-Spc containing 20 µM IPTG and grown to OD_600_ ∼0.5. Live cells were immobilized on agarose pads for microscopy. (ii) Colocalization of ZapA-mCh and MltA-mNG in Fig. 2D. A starter culture of EC5868 was grown overnight at 30°C in LB-Spc-Kan containing 10 µM IPTG. The culture was then diluted 1:200 into LB0N-Spc containing 20 µM IPTG and grown at 30°C to OD_600_ ∼0.5. Live cells were immobilized on agarose pads for microscopy. (iii) MltA-mCh in Fig. 3. “WT” refers to EC6447. A starter culture was grown overnight at 30°C in LB-Spc containing 10 µM IPTG. This culture was diluted 1:200 into LB-Spc containing 20 µM IPTG and grown at 30°C to OD_600_ ∼0.5, then back-diluted 1:10 into the same medium at either 30°C or 42°C and grown for 60 min. Live cells were immobilized on agarose pads for microscopy. “*ftsZ*(ts)” refers to EC6449. It was grown similarly except that the initial dilution was 1:500 and the 1:10 back-dilution was into either LB-Spc 20 µM IPTG at 30°C (permissive) or LB0N-Spc 20 µM IPTG at 42°C (non-permissive). “*ftsQ*(ts)” refers to EC6366. It was grown as described for “WT” except that samples were taken for microscopy after 90 min of growth under permissive/non-permissive conditions. “*ftsI*(ts)” refers to EC6362. It was grown as described for “WT” except that the non-permissive condition was LB-Spc and samples were taken for microscopy after 90 min of growth under permissive/non-permissive conditions. “*P_BAD_::ftsN*” refers to EC6681. An overnight culture was grown at 30°C in LB-Spc containing 20 µM IPTG and 0.2% Ara. In the morning this culture was diluted 1:50 into the same medium but with 0.2% glucose and grown at 30°C for 2 hours to OD_600_ ∼0.5. Then cells were washed and transferred into LB0N by centrifuging 1.0 ml of culture in a microfuge tube and taking up the resulting cell pellet in 1.0 ml of LB0N. This cell suspension was diluted 1:25 into LB0N-Spc containing 20 µM IPTG and either 0.5% Ara (permissive) or 0.2% Glu (non-permissive). Live cells were immobilized for microscopy when the cultures reached OD600 ∼0.3 (about 2 hours).

### Growth curves and morphology

Overnight cultures of EC251 (WT) or EC3546 (Δ*mltA*) that had been grown at 30°C in LB were diluted 1:20,000 into 25 ml of LB0N in a 250 ml Erlenmeyer baffled culture flask. The calculated starting OD_600_ was ∼0.00025. Cultures were incubated at 37°C and 210 rpm to stationary phase with periodic removal of 1 ml samples for OD_600_ measurement. At OD_600_ ∼0.5, cells were fixed by the addition of paraformaldehyde/glutalaldehyde-NaPO_4_ to final concentrations of 2.5%/0.8% and 20 mM, respectively. Fixed cells were stained with FM4-64 and immobilized on agarose pads for microscopy.

### Deoxycholate sensitivity

To test sensitivity to deoxycholate (DC) in spot titer assays, starter cultures were grown overnight at 37°C in LB or LB-Spc in the case of plasmid strains. Cells from 1 ml of culture were pelleted in a microcentrifuge, resuspended in LB0N and adjusted to a OD_600_ = 1.0 (defined as 10^0^). Then 10-fold serial dilutions (-1 to -6) were made in LB0N and 3-μl aliquots were spotted onto an LB0N agar plate containing 20 mM IPTG with or without 0.1% deoxycholate. Plates were photographed after 18 h at 37°C.

### Localization of GFP-FtsN^SPOR^ *in vivo*

Starter cultures of EC6534 and EC6539 grown overnight at 30°C in LB-Amp. The next day, these were diluted 1:200 into LB0N-Amp and grown at 30°C to OD_600_ ∼0.5. Live cells were spotted onto agarose pads for microscopy. It was not necessary to add IPTG to achieve adequate expression of *gfp-ftsN*^SPOR^ from the plasmid.

### Generation of anti-serum against MltA

Polyclonal anti-serum against *E. coli* MltA was raised in rabbits by ProSci Incorporated (Poway, CA). The protein used as antigen was obtained by constructing a pET vector encoding an N-terminal hexahistine tag to MltA (residues 22-359). The resulting plasmid, pDSW2409, was transformed into BL21(DE3). Cultures for protein overproduction were grown in LB-Amp at 37°C to OD_600_ = 0.5, induced by adding IPTG to 1 mM final concentration and grown for an additional 3 hours at 22 °C. Purification was by cobalt affinity chromatography according to instructions from the manufacturer (Takara, San Jose, CA). Purified His_6_-MltA(22-359) was dialyzed into 50 mM NaPO_4_ (pH 7.5), 150 mM NaCl. The preparation was >95% pure.

### Purification of AmiD, RlpA, GFP-DamX^SPOR^ and GFP-MltA(D328A)

Purification of hexahistidine-tagged *E. coli* AmiD, *P. aeruginosa* RlpA and *E. coli* GFP-DamX^SPOR^ proteins has been described (17). GFP-MltA(D328A) refers to a catalytically inactive mutant protein spanning codons 22-359, i.e., lacking the N-terminal signal sequence for export and lipid modification. This protein was overproduced in BL21(DE3) and purified as described above for the wild-type variant used to raise anti-sera against MltA. Purified His_6_-MltA(D328A) was dialyzed into 50 mM NaPO_4_ (pH 6.5), 200 mM NaCl, 5% glycerol and stored at -80 °C.

### Microscopy-based assay for binding of GFP-tagged proteins to purified PG sacculi

Sacculi from WT, Δ*mltA* or Δ*LTs E. coli* were purified as described (17). The corresponding strains are EC251, EC3546 and EC3708, respectively. Sacculi were immobilized on poly-L-lysine treated multiwell microscope slides and the binding assay was performed as described (17). Proteins were diluted from stock solutions into phosphate-buffered saline containing bovine serum albumin (PBS/BSA) and added to wells after the blocking step—either 10 µl of GFP-DamX^SPOR^ at 10 nM or 10 µl of GFP-MltA(D328A) at 100 nM. GFP-MltA(D328A) was used at higher concentration because it bound weakly on sacculi, consistent with it being an enzyme that must turn over substrate for efficient catalysis. Incubation was continued for 30 min before unbound protein was removed by washing the wells five times with 10 μL of PBS. Finally, 3 μl of PBS was added per well, a coverslip was mounted, and samples were examined immediately under the microscope.

To increase the abundance of denuded PG glycans, sacculi were treated with 10 μL of 2.5 µM AmiD. To decrease the abundance of denuded PG glycan while also increasing the abundance of 1,6-anhMurNAc ends, sacculi were treated with 10 µl of 5 µM *Pa*RlpA. Both proteins were in PBS/BSA and added after the blocking step. Slides were placed in a covered Petri dish containing droplets of water to minimize evaporation. Incubation was for 30 min at 37°C. Enzymes were removed by washing the wells five times with 10 μL of PBS, at which point 10 μL of GFP-DamX^SPOR^ or GFP-MltA(D328A) protein in BSA/PBS was added as above. Subsequent procedures were as described above.

### Image analysis and figure preparation

Cell length and septal localization of fluorescent proteins were measured and scored manually with cellSense Dimension software from at least 100 cells in at least two experiments. Fluorescence intensity across the midcell was quantified in Fig. 4E using the Line Profile tool in ImageJ from three experiments. For this, ≥15 cells that exhibited septal localization were chosen at random, and their fluorescence profiles were captured. These were averaged. Fluorescence intensities varied from day to day, so it was not informative to pool data from different days without first normalizing. This was done by setting the maximum pixel intensity for GFP-FtsN^SPOR^ in the Δ*mltA* background on any given day to 1. Pixel intensities for the wild-type background were normalized using the raw average intensity value for the Δ*mltA* background on that day. In other words, in each experimental replicate (different days), the raw average maximum pixel intensity for GFP-FtsN^SPOR^ was about 20% higher in the Δ*mltA* background than in wild-type.

Adobe Photoshop and Adobe Illustrator were used to adjust and crop images. Adjustments used the Levels, Brightness, and Contrast tools, and some images were inverted to improve legibility.

### Statistics

Microsoft Excel was used to perform two-tailed Student’s t-Tests.

## ACKNOWLEDGMENTS

This work was supported by NIH R01GM125656 (to D.S.W.) and in part by T32AI007511 to G.M.K. We thank Thomas Bernhardt for the AmiD overproduction plasmid and Matt Jorgenson for helpful discussions. DNA sequencing was performed at the Genomics Division of the Iowa Institute of Human Genetics, which is supported in part by the University of Iowa Carver College of Medicine.

